# Symmetry breaking during morphogenesis of a mechanosensory organ

**DOI:** 10.1101/718502

**Authors:** A. Erzberger, A. Jacobo, A. Dasgupta, A. J. Hudspeth

**Affiliations:** Howard Hughes Medical Institute and Laboratory of Sensory Neuroscience, The Rockefeller University, New York, NY 10065 USA

## Abstract

Actively regulated symmetry breaking, which is ubiquitous in biological cells, underlies phenomena such as directed cellular movement and morphological polarization. Here we investigate how an organ-level polarity pattern emerges through symmetry breaking at the cellular level during the formation of a mechanosensory organ. Combining theory, genetic perturbations, and *in vivo* imaging assisted by deep learning, we study the development and regeneration of the fluid-motion sensors in the zebrafish’s lateral line. We find that two interacting symmetry-breaking events — one mediated by biochemical signaling and the other by cellular mechanics — give rise to a novel form of collective cell migration, which produces a mirror-symmetric polarity pattern in the receptor organ.

## Introduction

Broken symmetries in living matter often stem from regulated biological processes across different spatiotemporal scales. During development or regeneration, for example, organ-scale structures and patterns emerge as a result of broken symmetries at the level of the constitutive cells [1–4]. Cellular polarity gives rise to systematic spatial asymmetries in features such as morphology, protein distribution, and mechanical properties [5–7].

Because of their highly stereotyped structure, the auditory, vestibular, and lateral-line sensory systems are especially attractive subjects for investigations of symmetry breaking in biological patterning. The mechanosensory epithelia of these organs contain precisely arranged sensory hair cells surrounded by non-sensory supporting cells. The top surface of each hair cell bears a mechanically sensitive organelle, the hair bundle, a cluster of rod-like processes that increase in length in one direction (Fig. 1A). This structural asymmetry accords with functional anisotropy: deflecting the bundle toward its tall edge excites the associated nerve fibers, whereas movement in the opposite direction has the converse effect [8, 9].

**Figure 1:**
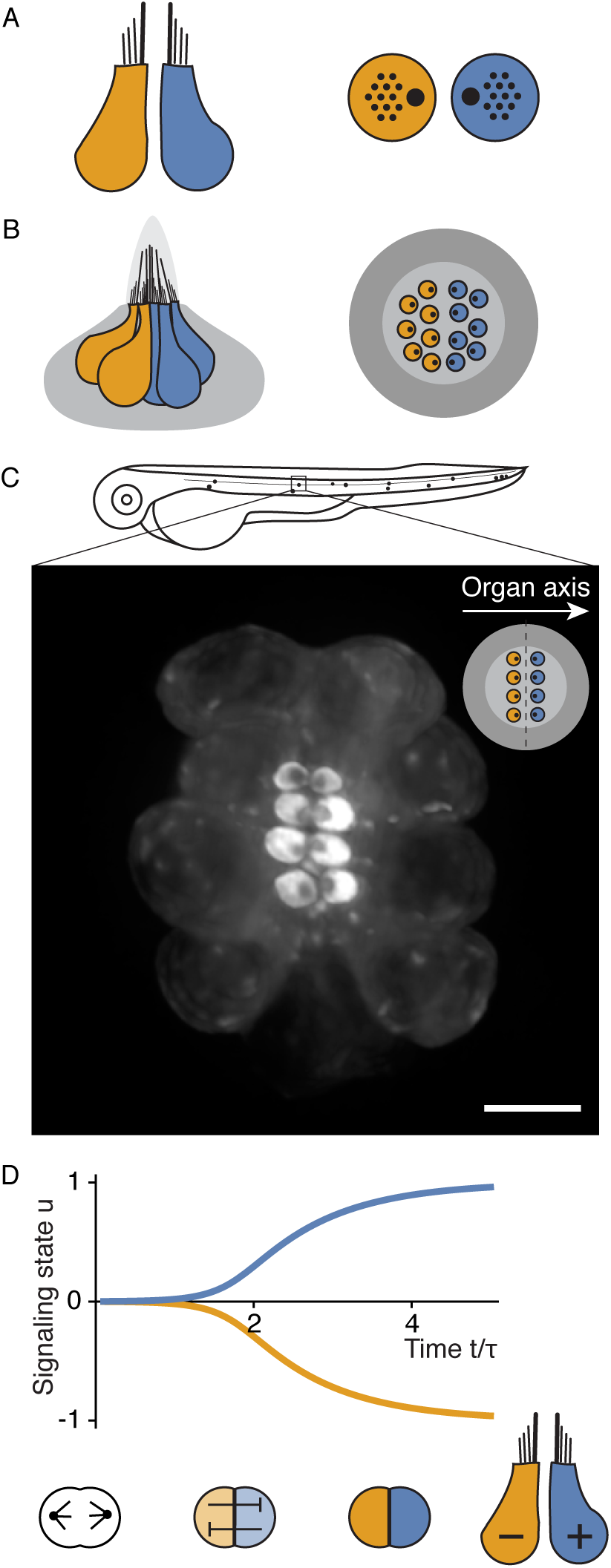
Multi-scale polarity in lateral-line neuromasts. **A**, Schematic side and top views portray a pair of oppositely polarized hair cells, the mechanoreceptors of the auditory, vestibular, and lateral-line systems. The asymmetric hair bundle protruding from the apical surface of each hair cell responds to mechanical deflection in a directionally sensitive manner. Many mechanosensory epithelia contain oppositely oriented hair cells, aligned to a global polarity axis; this arrangement permits a precise measurement of stimuli delivered along the organ axis. **B**, Polarized sensory hair cells are arranged in a mirror-symmetric pattern across a lateral-line neuromast of the larval zebrafish. Hair cells are surrounded by supporting cells (gray). **C**, As shown in the schematic diagram, neuromasts in the lateral-line organ of a zebrafish larva are arrayed along the anteroposterior body axis. The fluorescence micrograph of a neuromast from a transgenic zebrafish larva shows hair cells labeled with *β*-actin-GFP against the dark background of unlabeled supporting cells. The apical surfaces of the hair cells appear as bright discs with asymmetrically localized dark spots that reveal the orientations of the hair bundles. The inset shows this arrangement schematically. Scale bar: 5 µm. **D**, The hair cells of neuromasts are formed in pairs by the division of a precursor cell. The nascent hair cells subsequently engage in mutually suppressive intercellular signaling. A model of the signaling circuit illustrates the resulting bistability in the signaling state *u* of the cells: small initial differences between interacting cells are amplified over a timescale *τ* to yield opposite cellular polarities (Eq. 1; Supplementary Note 1).

In order to respond effectively to various stimuli, the hair bundles in each sensory organ must adopt a specific pattern of orientations. A molecular signaling system termed planar cell polarity (PCP) mediates cell-cell interactions that align the hair bundles of neighboring hair cells, rather like the nematic interactions between anisotropic particles. In many epithelia, long-range cues additionally modulate PCP signaling, promoting a state with uniform nematic ordering, in which all cells are aligned to a particular axis across an organ [10, 11].

The hair bundles in the human vestibular system as well as those in the lateral-line organs of many aquatic animals are arranged in a mirror-symmetric pattern with respect to the organ axis. This arrangement facilitates the precise measurement of stimuli delivered in opposite directions along a specific axis of sensitivity. To consider the example examined in this study, each side of the tail of a larval zebrafish bears discrete clusters, called neuromasts, with about 20 hair cells apiece. Although the PCP pattern is uniform across all cells in a neuromast and accords with the organ axis, the polarity of half of the hair bundles is reversed with respect to this axis (Fig. 1B,C; Supplementary Video 1; [12–14]). As a result, half of the hair bundles in a neuromast oriented along the anteroposterior body axis are sensitive to tailward water motions, which occur for example when the animal swims forward. The complementary half of the bundles respond to headward flows, which are particularly significant when a predator — usually a larger fish — strikes from behind.

Hair cells of the two orientations originate in pairs from the division of a precursor cell [13]. Rather than through an asymmetry in the division itself, the opposite orientations arise from a subsequent breaking of symmetry between the two daughter cells through a biochemical interaction [15]. Immediately after division, both nascent hair cells express similar amounts of the signaling protein Notch, but mutually inhibitory interactions create a bistable system in which the two cells diverge to distinct states. These interactions occur through a direct cell-cell contact and result in the accumulation of the Notch intracellular domain (NICD) in one of the cells and its depletion in the other. Describing the signaling state of each cell *i* with a unitless variable *u*_*i*_, the bistability is captured by the following minimal model

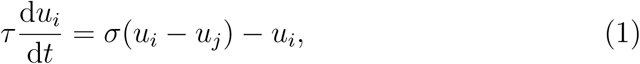

with *σ*(*u*) a generic sigmoidal function (Fig. 1D; Supplementary Note 1; [16]). Any small asymmetry in the initial conditions amplifies over the timescale *τ* and gives rise to a positive and a negative state. The NICD-positive and -negative sibling cells subsequently develop opposite polarities [17, 15].

To investigate the spatial arrangement of the cells, it is helpful to introduce the signaling dipole

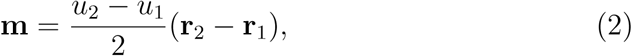

in which **r**_*i*_ denotes the position of cell *i*. In reference to the PCP axis, a positive dipole corresponds to a pair whose mature hair bundles are apposed at their tall edges, whereas in a negative dipole the tall edges of the two hair bundles face away from one another. Although *a priori* the signaling-induced symmetry breaking between the daughters can give rise to both positive and negative dipoles at equal probability, mature hair-cell pairs are exclusively found in positive-dipole configurations, precisely aligned with the organ axis both during normal development and when regenerating in response to damage (Supplementary Fig. 1; Supplementary Note 1; [13]).

Charge dipoles align in an external electric field; but how do pairs of hair cells organize in space to give rise to the observed organ-scale broken symmetry? We hypothesize that, rather than being driven by external input, the biological pattern emerges from cell-level interactions. This is suggested in particular by the fact that neuromasts display remarkable morphogenetic robustness. In contrast to mammalian mechanosensory organs, in which the loss of hair cells leads to irreparable disorders of hearing and balance, neuromasts possess an inexhaustible regenerative capacity: fully functional and correctly patterned neuromasts regenerate after virtually any extent of damage [18, 19]. Moreover, the large-scale organ patterns are maintained in a highly dynamic environment, because neuromast cells undergo continuous turnover throughout the lifetime of the organism [20]. Here we show that a combination of symmetry-breaking events mediated by biochemical signaling and active mechanical forces at the cellular level establish an organ-scale symmetry with a pattern of oppositely oriented hair bundles. Furthermore, our results suggest a robust mechanism for the precise mosaic arrangement of hair cells and supporting cells.

## Results

### Dynamics of contact angles during differentiation

How does the precise spatial arrangement of cells in the neuromast arise? To approach this issue, we studied the shape changes, movements, and mechanical properties of maturing hair cells over time. We began by investigating the changes in the cell-cell contacts over the course of maturation. During early stages of their development, sibling hair cells engage in Notch signaling across a direct cellular interface, but mature and functional hair cells are fully surrounded by glia-like supporting cells (Fig. 1B,C; [21]). The cells must therefore undergo a substantial reorganization of their contact geometry during maturation.

By expressing fluorescent marker proteins under the control of specific promoters, we labeled individual hair cells and visualized them against the dark background of the unlabeled supporting cells (Figs. 1C and 2A; Supplementary Video 1). We observed that, after precursor division, the daughter cells formed a flat interface with a contact angle *α* ≈ 90° and remained in that configuration for several hours (Fig. 2B). The contact angle sub-sequently decreased and the cells detached from one another, allowing the intercalation of supporting cells. Using deep learning-assisted live imaging to minimize photo-oxidative damage from protracted microscopy, we quantified the contact angle between hair-cell pairs from progenitor division until cell-cell detachment and documented highly stereotyped dynamics across 24 hair-cell pairs from different neuromasts (Fig. 2C; Supplementary Video 2).

**Figure 2:**
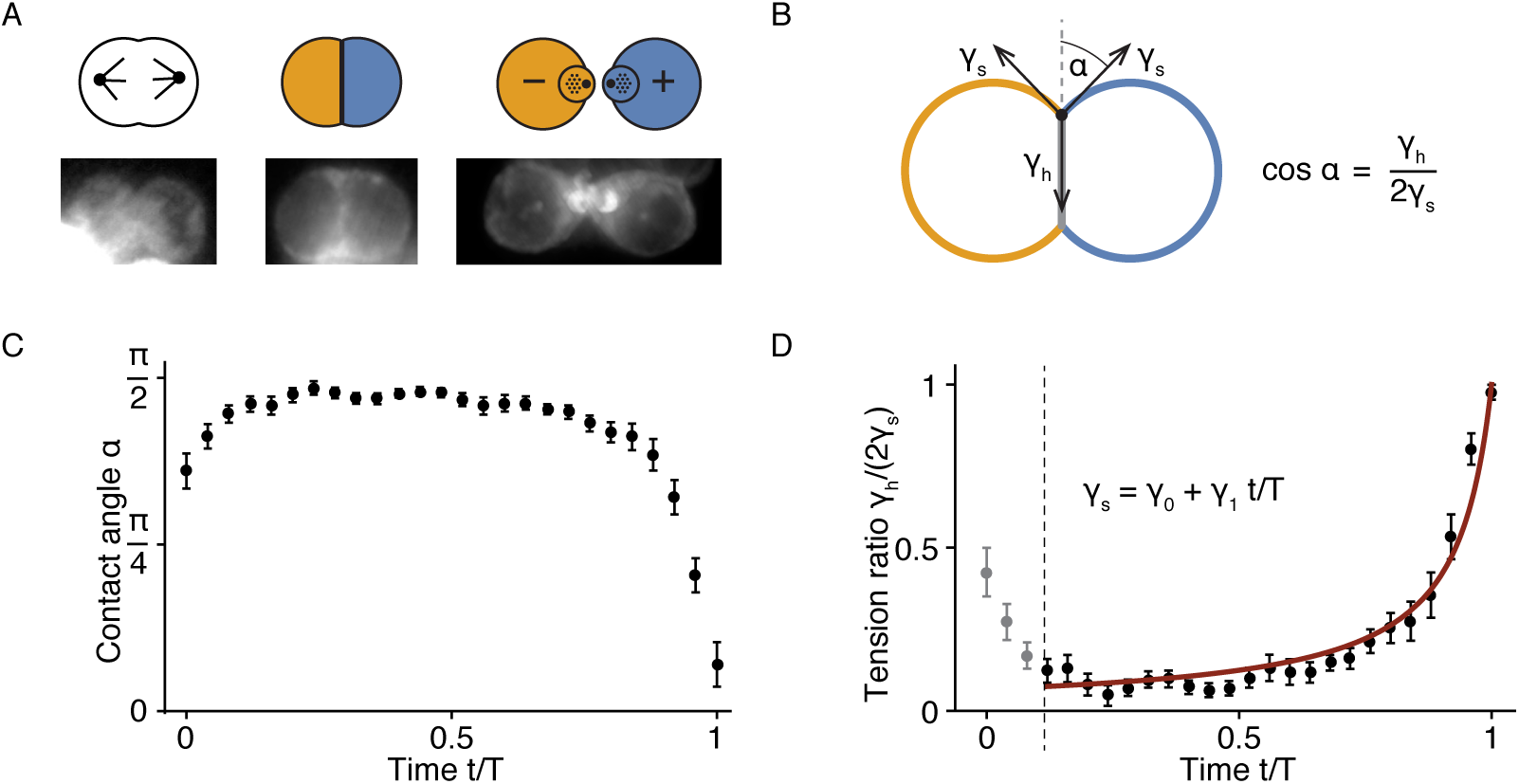
Contact-angle dynamics of maturing hair-cell pairs. **A**, Neuromast hair cells undergo a stereotypical series of shape changes during maturation. Following division, they form a round doublet with a flattened contact surface before developing apical surfaces with hair bundles. Mature hair cells are always completely surrounded by supporting cells. The lower panels show fluorescent images of the corresponding stages from larvae expressing *β*-actin-GFP in hair cells. **B**, The contact angle *α* between the cells is controlled by the cellular surface tensions *γ*_h_ and *γ*_s_, in which *γ*_h_ is associated with the interface between the two hair cells and *γ*_s_ with that between hair cells and supporting cells (see Supplementary Note 2.2). **C**, As shown on a timescale normalized to the doublet lifetime *T*, the contact angles in 24 pairs of developing hair cells underwent stereotyped dynamics between cell division and cell-cell detachment. After mitosis, each cell doublet assumed a round shape that persisted for several hours. The contact angle then decreased sharply until the cells detached from each other. **D**, The timecourse of the ratio of surface tensions for the data in panel **C** accords with a first-order linear decrease in *γ*_s_ with a single fitting parameter *γ*_1_*/γ*_0_ = *-*0.93 ± 0.01 (see Supplementary Note 2.2). The initial decrease in the surface tension ratio likely reflects an overall loss of cell contractility following mitotic rounding. The error bars denote SEMs.

To obtain insight into the mechanical processes underlying changes in the intercellular interfaces, we considered the associated effective surface tensions. The cellular surface tension is a mesoscopic variable arising from a variety of active and passive molecular processes, including intercellular adhesion at the surface membranes and active contractility of the underlying actomyosin cytoskeleton [22–25]. The surface tension *γ* can be defined as an effective energetic cost per unit area for the associated interface: it is conjugate to the interfacial area *𝒜* in the expression for the surface potential of the cell doublet [26]. A hair-cell doublet has two types of interfaces: the contact area between the two hair cells *𝒜*_h_, and the contacts with the surrounding supporting cells *𝒜*_s_. Accordingly, the effective surface energy is given by

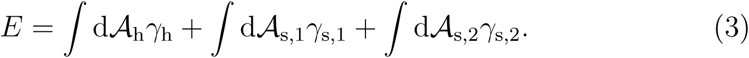

When the cell doublet is in a configuration that minimizes this energy, the contact angle *α* between the two cells is controlled by the surface tensions (Fig. 2B; Supplementary Fig. 2; Supplementary Note 2.1). If the supporting cell tensions in the contact point are equal *γ*_s,*i*_ = *γ*_s_, the contact angle is

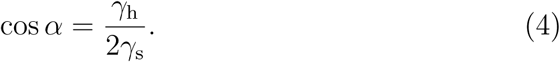

We used Eq. 4 to infer the ratio of the interfacial tensions from our measurements of the contact angle over time (Fig. 2D). Cellular surface tension is subject to active spatiotemporal regulation: various biochemical and genetic processes modulate the adhesive and contractile properties of cells. Indeed, hair cells and supporting cells in the mammalian cochlea differentially express heterophilic adhesion molecules whose interactions underlie the mosaic arrangement of the cells required for sensory function [27, 28]. We inquired whether differential adhesion might explain the characteristic contact dynamics of nascent hair cells in a neuromast. If adhesion molecules that mediate heterophilic interactions with supporting cells are expressed under a hair cell-specific genetic program, the expected mesoscale effect is a gradual decrease of *γ*_s_ during differentiation. Indeed, we found that the dynamics of the contact angle is reproduced surprisingly well by a single-parameter fit of a first-order decline in *γ*_s_ over time (Fig. 2D). This finding suggested that the morphological rearrangements were driven primarily by changes in the strength of the adhesive interactions between hair cells and supporting cells.

### Polarity of active protrusions and directed cellular movements

During the later stages of their maturation, hair cells extend stout cytoskeletal protrusions that resemble the actin-filled lamellipodia and filopodia of migrating cells [29, 21]. Within the context of surface tension, such protrusions can be interpreted as the signature of a “wetting” transition: the interfacial tension *γ*_s,*i*_ crosses a threshold beyond which the corresponding interfacial area is maximized rather than minimized under the constraints of the system [30, 31]. We observed that, rather than being uniformly distributed, protrusions were restricted to particular cellular regions: the hair cells of each pair formed protrusions predominantly at their opposite poles (Fig. 3A; Supplementary Video 2). This observation suggested that, rather than decreasing uniformly over time, the interfacial tension *γ*_s,*i*_ also developed a spatial broken symmetry. A cellular surface that is subject to non-uniform tension may become protrusive in spatially restricted regions where the tension is locally below the wetting threshold. In particular, oriented gradients of surface tension may produce directed protrusive activity along a particular cellular axis [32–34].

**Figure 3:**
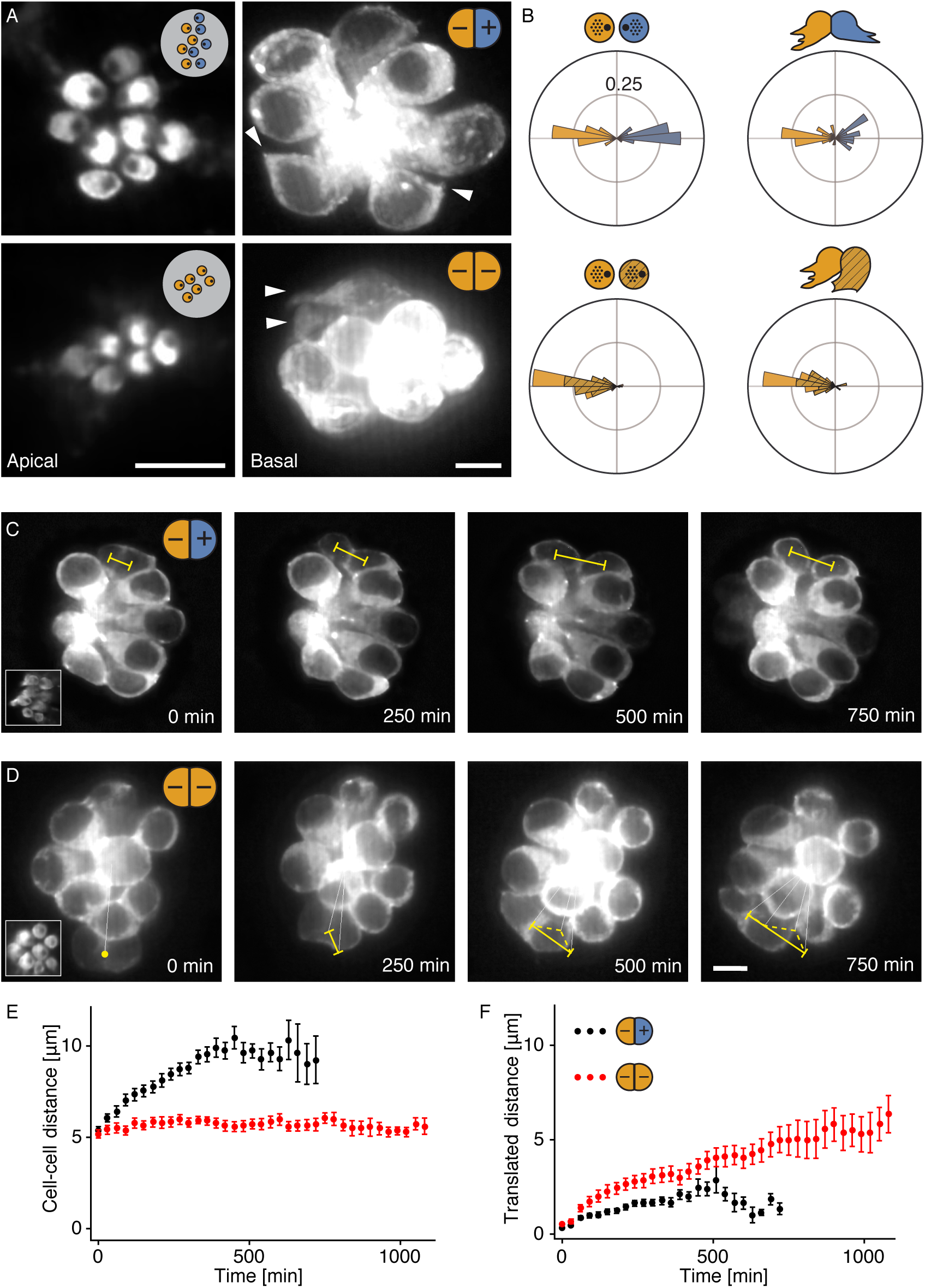
Relationship of polarity to cellular protrusions and movements. **A**, Fluorescence micrographs show *β*-actin-GFP in a neuromast of a wild-type larva (top) and a mutant lacking functional Notch protein (bottom). The hair bundles on the apical surface (left) are oppositely oriented in the wild-type animal, but those in the Notch mutant are uniformly oriented. In the mutant larvae the bistable signaling system that regulates polarity reversal is disrupted, so both sibling cells adopt the same polarity fate. The maturing hair cells in the neuromasts form distinct actin protrusions (right). The hair cells in the oppositely polarized pair of the wild-type extend oppositely oriented protrusions, whereas the cells in the Notch mutant extend protrusions in the same direction (arrowheads). Here and in panels C and D, insets in the first images reveal the polarities of the hair cells. **B**, Angular histograms show the orientations of hair bundles (left) and of cellular protrusions (right) relative to the axis of PCP in hair-cell pairs from wild-type larvae (top) and Notch mutants (bottom). **C**, In oppositely polarized pairs of hair cells, the center-to-center distance between cells increases during the protrusive stage of maturation, but net translation within the neuromast is minimal. **D**, Instead of moving away from each other, the uniformly polarized hair cells in Notch mutants translate as pairs in the direction of protrusive activity. **E**, The average timecourse for the intercellular separation of 28 hair-cell pairs from wild-type larvae (black) differs from that of 31 pairs from Notch mutants (red). **F**, The data for mean translated distance show that wild-type cells (black) separate almost symmetrically from their starting positions, whereas mutant cells (red) undergo a net migration. The error bars denote SEMs. Scale bars: 5 µm.

To establish whether the localized protrusions were directed by the same signaling system as that determining the orientation of hair bundles, we measured both the orientations of the protrusions and the angles of hairbundle polarity relative to the axis of PCP. In pairs of oppositely polarized hair cells, the orientations of protrusions were indeed closely aligned with the polarities of the hair bundles (Fig. 3B; Supplementary Video 2). We also imaged mutant fish that lacked functional Notch receptors so that the chemical circuit regulating polarity reversal was disrupted. The neuromasts of these mutants had uniformly polarized hair cells [15]. We found that the protrusions in the immature hair cells of the Notch mutants were also uniformly oriented in the direction defined by the orientations of the hair bundles (Fig. 3B; Supplementary Video 3).

Pairs of cells with oppositely polarized hair bundles occasionally occurred in Notch mutants, possibly as a result of compensatory effects from redundancies in the regulatory circuit. In these pairs, the cytoskeletal organization and protrusions were indistinguishable from those in wild-type animals (Supplementary Fig. 3). The misorientation of protrusions in uniformly polarized pairs was therefore not an effect of a systemic disruption owing to the mutation, but stemmed specifically from a lack of distinct polarities in these hair cells.

If polarized protrusions represent a signature of a broken symmetry in cellular surface tension — or more generally in the distribution of local forces exerted by the cell on its environment — might polarized forces produce active cellular movements? By monitoring the protrusive phase in both wild-type and Notch mutant neuromasts, we ascertained that hair cells in both conditions moved actively in the direction of their protrusions. In oppositely polarized pairs of hair cells, these movements increased the separation between daughter cells — a process that might facilitate the interposition of supporting cells — while the centroid of the pair remained stationary (Fig. 3C). By contrast, in uniformly polarized mutant pairs the intercellular separation remained small as the cells translocated together in the direction of their protrusions (Fig. 3D). In both instances, polarity-associated cellular movements appeared to be constrained eventually by the mechanical anchor-ages at the apical cellular surfaces (Fig. 3E,F).

We additionally investigated mutant neuromasts in which hair cells of both polarities arose from the experimentally induced expression of NICD in a random pattern rather than through intercellular interactions [15]. Even though positively and negatively polarized cells originated at random positions within these neuromasts, we predicted that polarity-based active movements would sort the cells into the appropriate halves of the neuromast and thus rescue bilateral symmetry. Our measurements of the hair-cell positions along the neuromast axes in wild-type and mutant animals confirmed this prediction (Supplementary Fig. 4). Taken together, our results suggest that hair-cell polarity not only defines the axis of hair-bundle orientation and the corresponding direction of maximal mechanical sensitivity, but also controls the directed protrusive and motile activities in maturing hair cells.

**Figure 4:**
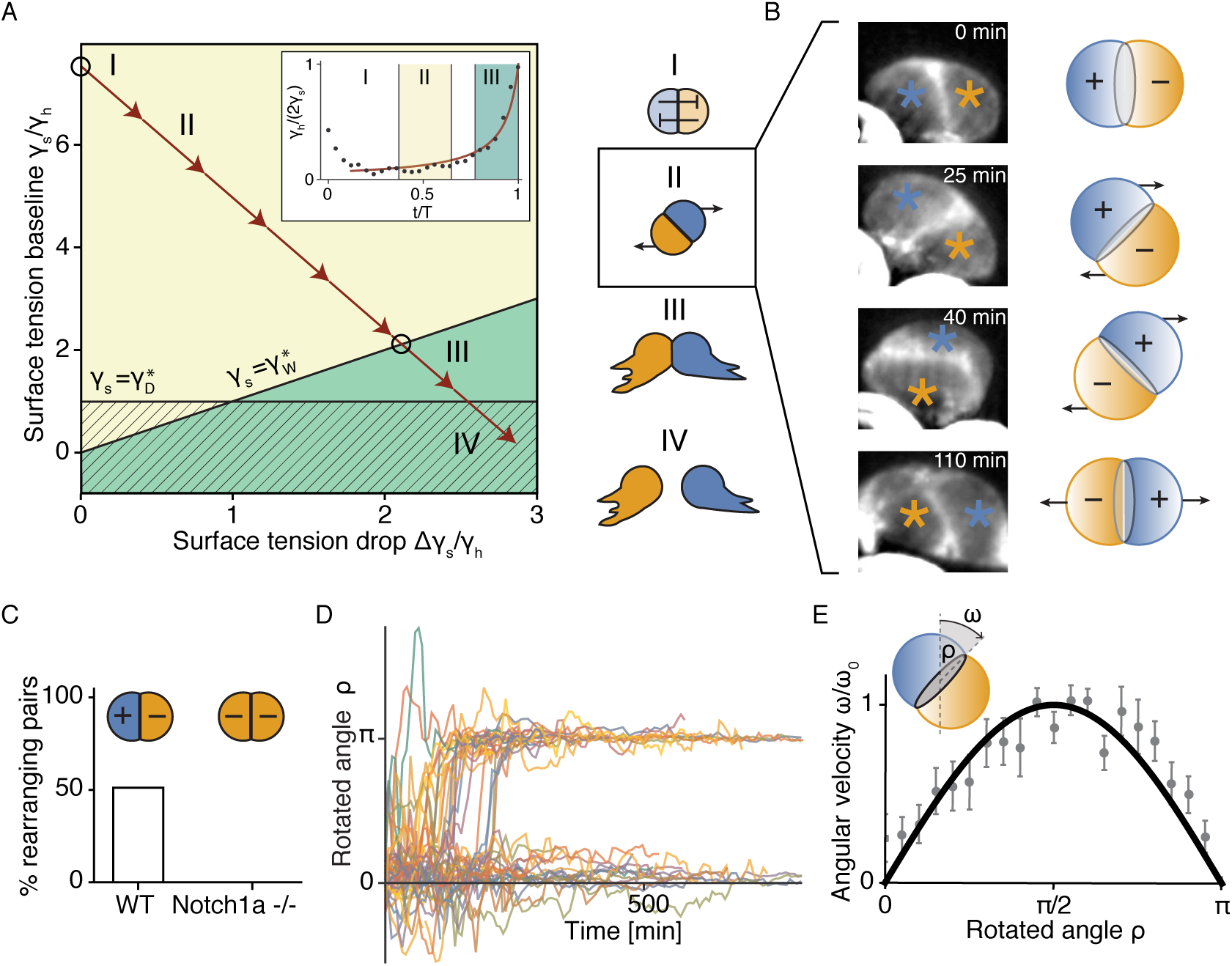
Surface mechanics of hair-cell maturation and active dipole transitions. **A**, A state diagram depicts the possible behaviors and morphologies of developing hair-cell pairs in the presence of a polarity-induced broken symmetry in the surface tensions *γ*_s,*i*_ (Eq. 5). The distinct regimes of the baseline tension *γ*_s_ and the tension drop Δ*γ*_s_ are bounded by the two critical lines 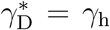 and 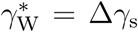. The detachment threshold 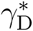 indicates where the interface between the hair cells disappears, and Min[*γ*_s,*i*_(*x−x*_0_)] = 0 at 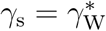, for which the “wetting” threshold is crossed at the cellular poles. The two cells form actin protrusions within the green region and are wholly separated by supporting cells within the hatched region. For Δ*γ*_s_ *>* 0, cell pairs can undergo dipole transitions from negative to positive configurations (yellow region). The approximate state-space trajectory of the cell doublet over the course of maturation (red arrows) is mapped by measuring the timing of hair-cell maturation events relative to our estimate of the dynamics of the surface-tension ratio (inset; Fig. 2). In particular, determining the onset of actin protrusions allows us to pinpoint the combination of parameters at the wetting transition (black circle). The successive stages of hair-cell maturation in the schematic diagrams include Notch signaling (I), dipole transitions (II), protrusion formation (III), and cell-cell detachment (IV). **B**, Dipole transitions occured in a fraction of cell pairs, in which the daughter cells switched position along the axis of PCP during the early stages of maturation. Fluorescent images from a cell pair expressing *β*-actin-GFP demonstrate the nearly round shape of the cell pair during these rearrangements. **C**, Twenty-two of 43 hair-cell pairs underwent a dipole transition in wild-type neuromasts with oppositely polarized hair-cell pairs, but none of the 18 uniformly polarized pairs in Notch mutants did so. **D**, The angular trajectories of the wild-type hair cells displayed sharply binary outcomes: the cells either completely switched position or remained stationary. **E** The angular velocity profile of the 22 rearranging pairs agrees quantitatively with the theoretical prediction *ω* = *ω*_0_ sin *ρ* with a single fitting parameter *ω*_0_ = (0.25 ± 0.01)10^*-*3^*·π/s*. The error bars denote SEMs.

### Coordination of hair-cell maturation by surface mechanics

We next explored the theoretical implications of a broken symmetry in the hair cell’s mechanical properties. In particular, our observations suggested a spatial variation of the surface tensions associated with the supporting-cell interfaces along the axis of polarity. We considered small perturbations from the uniform case which weakly depend on a cell-intrinsic coordinate (*x-x*_*i*_), in which *x* is the global organ axis and *x*_*i*_ is a reference point within cell *i*. With Δ*γ*_s_ denoting the magnitude of the tension drop along the cell, we considered the following surface tension profile for cell *i*:

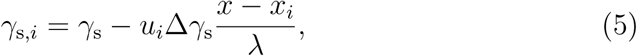

in which *λ* is a gradient length scale of the order of a cell’s size. The sign of the gradient along *x* is given by the signaling states *u*_*i*_, such that oppositely polarized cells have oppositely oriented surface-tension gradients (Fig. 1D).

The state space of the cell pair for different values of *γ*_s_ and Δ*γ*_s_ shows the critical line of the transition, at which the protrusions appear first at the cellular poles (Fig. 4A). To infer the approximate course of the cell pair through the state space during maturation, we estimated the overall decline in *γ*_s_*/γ*_h_ from our measurements of the contact-angle dynamics (Figs. 2C and 4A). Although we have no direct readout for the dynamics of the tension drop, we could establish the initial point by assuming that Δ*γ*_s_ is zero during the signaling phase, when the cells have not yet committed to a polarity fate. By measuring the time when the first protrusion appeared on the surface of each cell, and relating the corresponding contact angle to the surface-tension ratio, we additionally established when the cell doublet crossed the effective wetting threshold.

The interpolated state-space trajectory passes through three distinct regimes, which correspond to the observed successive stages of hair-cell maturation: the initial round morphology, the formation of protrusions, and the separation of daughter cells (Supplementary Video 2). Only two surface-mechanical changes are therefore required to explain the precise sequence of hair-cell maturation events: first, a differentiation-induced increase in heterophilic adhesion leading to a decline in *γ*_s_; and second, a polarity signaling-induced increase in Δ*γ*_s_. Note that these two processes are themselves coupled: the large cell-cell interface in the initial configuration *enables* the signaling interaction, which in turn *induces* the spatial broken symmetry that appears in the formation of polarized protrusions and concludes in the detachment of the two cells. In summary, a highly parsimonious theoretical framework self-consistently explains the key phenomena of hair-cell maturation.

### Polarity-driven transitions from negative to positive cell-pair dipoles

The polarity-directed broken symmetry in the surface tension also allows cell pairs to actively transition from negative-to positive-dipole configurations during maturation. In fact, such rearrangements of nascent pairs of hair cells have been observed in lateral-line neuromasts (Fig. 4B; Supplementary Video 4; [35, 36]). We quantified the trajectories of 43 nascent hair-cell pairs over time and observed that 22 underwent a dipole transition, whereas the remainder retained their initial configuration. In contrast, none of the 18 uniformly polarized hair-cell pairs that we observed in Notch mutants underwent a rearrangement (Fig. 4C–D). We did however observe a rearrangement in one of the oppositely polarized pairs that rarely occurred in the mutant background (Supplementary Fig. 3). This observation further supports the inference that the rearrangements are driven by the opposing polarities in a pair.

Dipole transitions consistently occurred during the initial stages of maturation, and took place over a period of up to 100 min, during which the contact angle between the cells was nearly constant and close to a value of 90° (Fig. 4A–B). This simple geometry permitted an analytical treatment of the surface mechanics and a prediction for the angular velocity *ω* of the dipole as a function of the rotated angle *ρ* to linear order in perturbations around an initial reference state for which *γ*_s,*i*_ = *γ*_s_. By calculating the movement of the contact line between the cells in response to the difference in surface tensions acting on it (Supplementary Note 2.4), we obtained

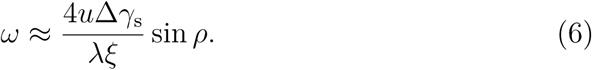

The angular velocity of a cell pair dipole thus depends on the magnitude of the signaling state *u* = |*u*_*i*_|, the surface tension drop Δ*γ*_s_*/λ* and a generic dissipative coeffient *ξ*. Fitting a single constant of proportionality *ω*_0_ to the measured average velocity profile produced a good agreement between theory and data (Fig. 4E).

### Interacting symmetry-breaking events

The biochemical and mechanical mechanisms underlying dipole formation can be described by a unified theory that elucidates the principle of polarity patterning in this system. The initial symmetry breaking between the two cells is triggered by bistable chemical signaling, and gives rise to positive and negative signaling states *u*_1_ = *-u*_2_ (Eq. 1). This process can be represented as the relaxation of the dipole **m** (Eq. 2) within an energy landscape that resembles a rotationally symmetric Goldstone potential (Fig. 5A; Supplementary Note 1). Immediately after division of a precursor, the pair of nascent hair cells occupies the unstable state atop the central peak. The system then relaxes to one of the degenerate minima around the base, and the polarities of the two cells are established. *A priori*, the potential is invariant with respect to the dipole angle *ψ*, but the subsequent manifestation of cellular polarity in the form of oppositely oriented forces gives rise to a second explicit symmetry breaking: the appearance of polarity-induced gradients in surface tension tilts the potential landscape along the global axis of the organ. This process eliminates the degeneracy of the minima at the bottom of the well and yields a unique stable state for the cell pair. The set of states at the bottom of the tilted potential corresponds to the effective surface potential of the cell doublet (Eq. 3), in which the dependence on the organ axis *x* appears explicitly through Eqs. 5. The surface-tension gradients are initially small, so the tilt can be considered linear, such that the surface potential can be simplified to

**Figure 5:**
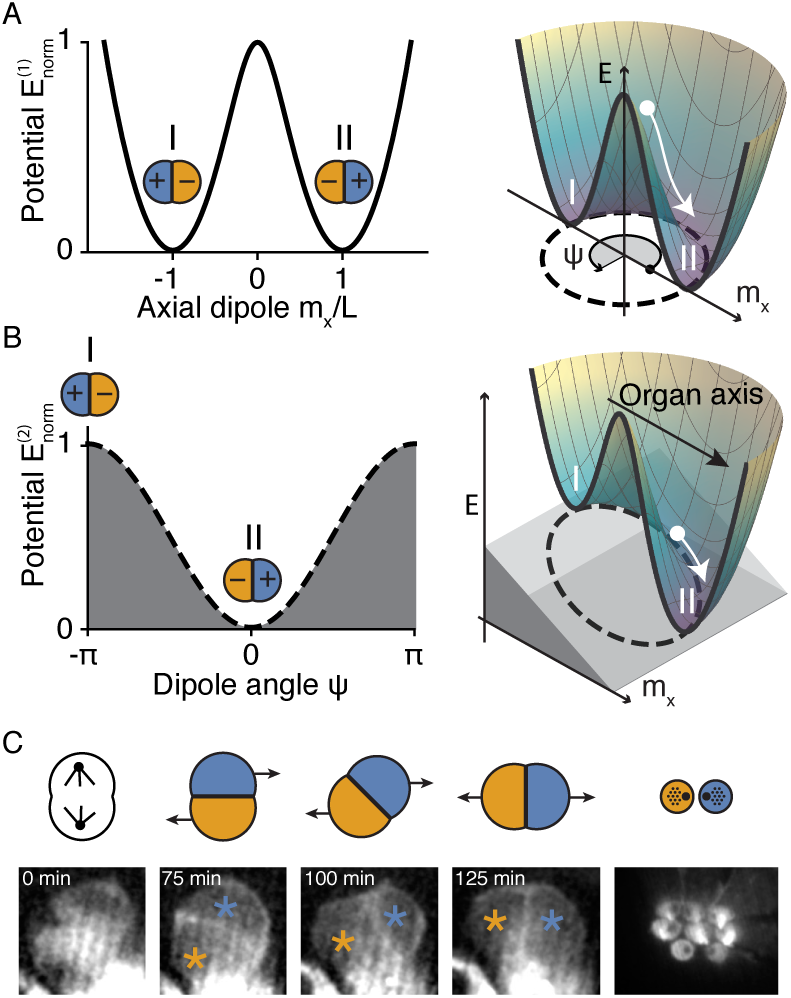
Symmetry-breaking events underlying polarity patterning. **A**, Biochemical signaling breaks the symmetry between the two sibling cells, creating a non-zero pair dipole **m**. The mutually suppressive Notch interactions give rise to a Goldstone-type signaling potential, as shown in a cross-sectional view (left) and portrayed in three dimensions (right). Initially similar cells poised near **m** = **0** move to one of the degenerate states at the bottom of the well (dashed line) as they undergo signaling. Because divisions of hair-cell precursors are often oriented along the organ axis *x*, most pairs have initial angles *ψ* = *π* (I) or *ψ* = 0 (II). **B**, Active mechanical forces subsequently give rise to an explicit symmetry-breaking event. Once the two hair cells have converged to positive and negative signaling states, they manifest opposite cellular polarities that induce active cell movements in opposite directions along the organ axis, by which the symmetry with respect to *ψ* is broken. The mechanical forces governed by cellular polarization tilt the Goldstone potential (right), such that the circular set of states at the bottom of the well assumes a cosine shape as a function of ψ (left), with a maximum at the negative and a minimum at the positive dipolar configuration (Eq. 7). **C**, The theory predicts that precursor cells dividing out of alignment with the axis of PCP should nonetheless produce hair-cell pairs that rearrange into the correct dipolar configuration. The products of occasional off-angle divisions indeed always attain the correct final state.

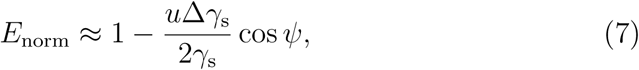

with *E*_norm_ = (*E -γ*_h_*𝒜*_h_)*/*(*γ*_s_*𝒜*_s,1_ + *γ*_s_*𝒜*_s,2_) (Supplementary Note 2.4.1). The unstable equilibrium positions at *ψ* = ±*π* correspond to the negative dipolar configuration of the pair, whereas the dipole is pointing along the organ axis in the minimum *ψ* = 0 (Fig. 5B). The derivative of the surface potential with respect to *ψ* defines the torque magnitude on **m** through

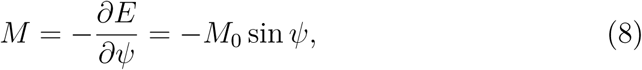

from which we can recover the angular velocity (Eq. 6) using the appropriate angle transformation.

The divisions of precursor cells in a neuromast are typically aligned with the organ axis as a consequence of PCP signaling, such that the initial distribution of cellular orientations is not uniform [13]. Rather than occupying all states at the bottom of the potential well with equal probability, the pairs cluster in two populations corresponding to positive and negativedipole configurations. When the explicit symmetry-breaking event renders the negative-dipole state unstable, the pairs at the top of the tilt transition to the stable positive-dipole configuration, whereas the remaining pairs maintain their original arrangement. An important prediction from our theory therefore concerns oblique cell divisions. For the proposed mechanism to produce the correct dipoles, PCP signaling is relevant for selecting the axis of the tilt (Fig. 5B), but not for aligning the axes of cell division. Once the potential landscape has been tilted, any pair of hair cells should transition to the unique minimum, irrespective of their initial orientation. As predicted, the occasional oblique divisions that we observed were always followed by partial rearrangements, precisely aligning the cell pair in the correct final configuration (Figs. 4D and 5C).

## Discussion

Combining experimental and theoretical approaches, we found that two interacting symmetry-breaking events mediate polarity patterning in the developing lateral-line system of the zebrafish. In the first event, bistable Notch signaling between the daughters of a cell division yields a pair of cells with positive and negative NICD states (Fig. 1; Supplementary Note 1). Consistent with the amplification of stochastic differences in the initial signal levels of the sibling cells after a symmetrical cell division, the signaling interaction produces both positive and negative-dipole configurations. However, a cell’s adoption of positive or negative NICD status influences not only the orientation of its hair bundle, but also the site at which cellular protrusions appear and the direction in which the cell migrates during maturation (Fig. 3). Oppositely oriented mechanical polarization thus constitutes a sub-sequent explicit symmetry breaking, which aligns the cell-pair dipoles and segregates the positive and negative cells to opposite halves of the organ (Figs. 4, 5). Note that these pairs behave like electric dipoles in an external electric field. In the biological system, however, rather than exerting a force on a charged particle, the external field merely selects an axis along which cell-intrinsic forces are exerted. These forces result from active processes within each cell’s cytoskeletal and adhesive machinery. Thus, cellular polarity drives *active* dipole transitions in this system.

Although surface tension-driven movements on substrates have been well explored both for passive droplets and for active cells [37, 38, 32], to our knowledge this study represents the first theoretical treatment of a system with two coupled, active “droplets” with oppositely oriented gradients in surface tension. The resulting phenomenon represents a novel form of collective motility, in which rather than moving uniformly—as in convergent extension [6]—the polarized cells undergo pairwise rearrangements that result in a mirror-symmetric polarity pattern at the organ scale. Relying on only two control parameters, our description of hair-cell surface mechanics not only quantitatively recapitulates the rearrangement trajectories and contact-angle dynamics measured *in vivo*, but also self-consistently explains the full sequence of distinct morphologies observed over the course of hair-cell maturation (Fig. 2; Fig. 4; Supplementary Note 2). In fact, the precise temporal regulation of the successive maturation events is parsimoniously interpreted as a natural consequence of two simple surface-mechanical changes in the hair cells. These mechanical processes are in turn triggered by two underlying differentiation events, first the commitment to become a hair cell, which introduces heterophilic interactions at the interface with the supporting cells; and subsequently the polarity-fate decision, which generates a spatial gradient in the surface tension.

The coupling of signaling and mechanics in maturing hair cells has an additional important effect: the two differentiation processes are themselves coordinated by their own geometrical manifestations. Although Notch signaling requires a direct contact surface between the interacting cells, mature and functional hair cells must be fully surrounded by supporting cells. Our findings imply that the system self-regulates the timing for this change in surface geometry. Because convergence to the positive or negative signaling state is a prerequisite for setting the cellular forces that ultimately separate the two cells, the cell-cell contact required for signaling infallably persists for as long as is required.

In conclusion, biochemical and mechanical interactions at the cellular level lead to the emergence of organ-scale patterns in neuromast morphogenesis. We propose that feedback effects coordinate the different cellular processes in time and promote robustness, suggesting that the regenerative capacity of this organ relies on these self-organizing principles. The coupling of directed mechanical forces during morphogenesis to functional and morphological cellular polarity in the mature stage provides a powerful mechanism for long-range patterning that bridges the levels of cells and organs.

## Materials and Methods

### 1 Animal handling and genetics

Experiments were performed in accordance with the standards of Rocke-feller University’s Institutional Animal Care and Use Committee. Zebrafish were raised in E3 medium (5 mM NaCl, 0.17 mM KCl, 0.33 mM CaCl_2_, 0.33 mM MgSO_4_, 1 µg*·*mL^*-*1^ methylene blue) in an incubator maintained at 28 ^°C^. Wild-type TL zebrafish were obtained from the Zebrafish International Resource Center. The following transgenic zebrafish lines were used: *Tg(myo6b:actb1-EGFP)* [39], *Tg(5xUAS-E1b:6xMYC-notch1a-intra)* [40], *Tg(myo6b:GAL4FF)* allele *ru1012Tg* [21], *and Tg(−8.0cldnb:lynEGFP)* [41]. In addition, we used the mutant line *Notch1a-/-* allele *b638* [42]. Embryos and larvae were staged as described [43].

#### 1.1 Treatments of larvae

To ablate mature hair cells in some experiments, we treated 3 dpf larvae for 2 hr at 28 ^°C^ with 1 µM CuSO_4_ in E3 medium. The animals were thoroughly washed and maintained overnight in E3 medium, then used for live imaging or fixed for immunofluorescence imaging on the following day.

### 2 Image acquisition and analysis

We performed time-lapse confocal fluorescence microscopy for up to 40 hrs on live animals. Embryos of 3–5 dpf were anesthetized in 600 µM 3-aminobenzoic acid ethyl ester methanesulfonate in E3 medium and mounted in a 35 mm glass-bottomed dish in 1% low-melting-point agarose.

Transgenic lines expressing *actb1-EGFP* were imaged under a 100X/NA 1.35 silicone-oil objective lens and an Olympus IX81 microscope equipped with a microlens-based, super-resolution confocal system (VT iSIM, VisiTech international). Neuromasts were imaged at intervals of 5 min as Z-stacks acquired with 0.5 µm steps under laser excitation at 488 nm. To keep the regions of interest within view over the imaging period, we developed a customized tracking algorithm that permits the automated updating of stage positions correcting for sample drift.

Transgenic lines expressing *lynEGFP* were imaged under a 60X/NA 1.3 silicone-oil objective lens with an Olympus IX81 microscope equipped with a spinning-disk confocal system (Ultraview, Perkin-Elmer, Waltham, MA). Neuromasts were imaged at intervals of 5 min as Z-stacks acquired with 0.5 µm steps under laser excitation at 488 nm.

Immunofluorescence labeling and imaging of wholemount larvae constitutively expressing NICD are described in [15].

#### 2.1 Deep learning-assisted long-term imaging

Because protrusions and movement at nascent stages as well as hair bundle orientation at mature stages had to be measured in the same cells, the data presented in Figs. 2–3 required long-term imaging at high temporal and spatial resolution. To prevent sample deterioration due to photodamage, we acquired images at low laser power and 300 ms exposures, then used a neural network to perform content-aware image restoration [44]. To train the network, we acquired both ground-truth images with high signal-to-noise ratios at high laser power, and images of the identical samples and imaging regions at low signal-to-noise ratios. The total size of the training dataset was 20 stacks (20 neuromasts) of 30 Z-slices each.

#### 2.2 Image analysis

Prior to image analysis, images were subjected to automatic drift (translation) correction with the MultiStackReg Fiji plugin [45] and cropping. For illustrative purposes, the images displayed in Figs. 3–5 were additionally processed by applying a 1000-point rolling-ball background subtraction algorithm and a three-point Gaussian blur filter.

Semi-automatic extraction of the quantities measured from time-lapse recordings of *Tg(myo6b:actb1-EGFP)* larvae (hair bundle polarities, contact angles, protrusion orientations, and cell trajectories) was performed with custom ImageJ (National Institutes of Health, Bethesda, MD) extensions written in Python.

The angular trajectories of hair-cell pairs in Fig. 4D were obtained as follows. In time-lapse recordings of neuromasts from *Tg(−8.0cldnb:lynEGFP)* larvae, the nascent hair-cell pair of interest was identified and the Z-slice near the centroid of the pair was manually selected for each frame. In this set of slices, the outlines of the two cells were segmented using a “morphological snakes” method [46]. After the algorithm had been initialized at the correct position by manual selection of points near the center of each cell on the first frame, it obtained automatically the outlines of the two cells and calculated their centroids, which were used to run the morphological segmentation algorithm in the subsequent frame. Occasional incorrect segmentations were resolved by manual re-selection of the cell centers. This algorithm produced a list of relative distances and angles as a function of time for each selected pair of hair cells. To compensate for variations in the imaging axis, we aligned the measured angles to the neuromast axis for each pair.

### 3 Calculations, data fitting, and visualization

Calculations shown in Supplementary Note 1–2, data visualization, data fitting (Figs. 2 and 4), and statistical testing was performed using Mathematica (Wolfram Mathematica 11.3.0.0). The average evolution of the contact angle shown in Fig. 2 was obtained as follows. We first measured the lifetime *T* of each cell-pair doublet by determining the timepoints of the cell division *t*_Div_ and cell-cell detachment *t*_Det_ events. We then measured the evolution of the contact angle over this period and collapsed the individual pair traces by normalizing the timepoints to the respective doublet lifetimes,

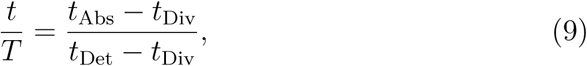

in which we have denoted the timestamps of the image recordings as *t*_Abs_. To average over all doublets, we computed linear interpolations between the datapoints for each doublet and averaged over evaluations taken at a resolution Δ*t/T* = 0.04.

For the average cell-cell trajectory data plotted in Fig. 3E, individual trajectories were aligned to the timepoint at which the first protrusion appeared without normalizing and averaged across pairs in 30 min bins. To obtain the angular velocity profile shown in Fig. 4D, the individual trajectories were first smoothed by applying an unweighted moving average filter with a 15 min window. The velocity was subsequently determined from each trajectory prior to averaging over the cell pairs.

## Acknowledgments

The authors thank A. Kaczynska for expert fish husbandry and A. Mietke, E. Siggia, and the members of our research group for critical reading and comments on the manuscript. A.E. was supported by a Feodor Lynen Fellowship from the Alexander von Humboldt Foundation and A.J. by an F.M. Kirby Postdoctoral Fellowship from Rockefeller University. A.D. is a Postdoctoral Associate and A.J.H. an Investigator of Howard Hughes Medical Institute.

## Author contributions

A.E. and A.J. designed the research with contributions from A.D. and A.J.H; A.J. performed the experiments with contributions from A.D; A.E. developed the theory; A.E. and A.J. analyzed the data; A.E. wrote the paper with contributions from A.J., A.D., and A.J.H.

