## Supplementary Materials for "Symmetry breaking during morphogenesis of a mechanosensory organ"

### Supplementary Materials:
### Symmetry breaking during morphogenesis of a mechanosensory organ

This document contains:

- Supplementary Video captions 1–4
- Supplementary Figures 1–5
- Supplementary Note
- Supplementary References

### Supplementary Video captions

**Supplementary Video 1: Sensory hair cells in a lateral line neuromast.** In a confocal recording enhanced by content-aware image restoration, a scan from the apex to the base of a neuromast in a four-day-old *Tg(myo6b:actb1-EGFP)* zebrafish larva shows sensory hair cells labeled with $\beta$-actin-GFP. The orientation of each hair bundle is revealed by the location of the kinocilium, which appears as a dark spot on the actin-enriched apical surface of each hair cell.

**Supplementary Video 2: Deep-learning-assisted, long-term imaging of hair-cell maturation.** A maximum-intensity projection of a time-lapse recording shows a developing neuromast in a *Tg(myo6b:actb1-EGFP)* zebrafish larva. We used content-aware image restoration (CARE; right) to improve the signal-to-noise ratio in images taken at low laser power and short exposure times (left). Minimizing photo-induced damage to the sample allowed us to image cell pairs throughout their maturation at high spatiotemporal resolution. Note the appearance of a new pair of hair cells in the lower left, followed by a second pair in the upper region. Each pair of daughter cells is initially rounded, with a flat contact surface between the cells. During a subsequent phase of protrusive activity the cells separate from one another. Finally, distinct apical surfaces with oppositely polarized hair bundles appear. In this and Supplementary Videos 3 and 4, the time signal is in hours and minutes.

**Supplementary Video 3: Hair-cell maturation in a Notch mutant larvae.** A CARE-processed maximum-intensity projection of a time-lapse recording depicts a developing neuromast in a *Tg(myo6b:actb1-EGFP)* animal lacking functional Notch1a receptors. The sensory hair-cell pairs form uniformly oriented actin protrusions during maturation and develop uniformly oriented hair bundles. Note the young pair of hair cells at the lower edge of the neuromast, both of which develop hair bundles oriented toward the animal's posterior. A second pair of nascent hair cells appears after four hours at the neuromast's upper margin.

**Supplementary Video 4: Dipole transition of a pair of nascent hair cells.** A CARE-enhanced time-lapse recording of a developing neuromast in a *Tg(myo6b:actb1-EGFP)* zebrafish larva is focussed at the level of hair-cell nuclei. The arrowhead in the first image denotes the location at which a progenitor cell appears and undergoes a division, giving rise to a pair of nascent hair cells. The two daughter cells maintain a flattened contact surface and undergo a rearrangement in which they switch positions along the PCP axis between 6:50 and 8:50. A second division near the upper edge of the neuromast yields a pair of cells that do not exchange places.

### Supplementary Figures

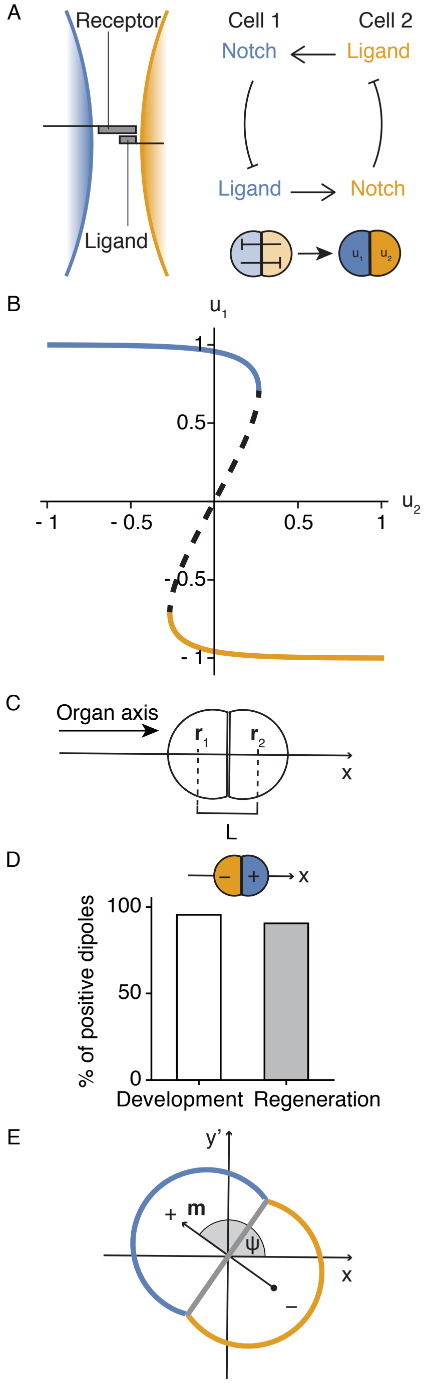


**Supplementary Figure 1: Notch signaling in hair cells and the signaling dipole. A,** Notch signaling relies on chemical interactions between receptor and ligand molecules. A ligand-receptor binding event across a cell-cell interface activates the receptor. In many contexts, receptor activation triggers the suppression of ligand expression in the same cell, resulting in a mutual inhibition circuit with bistable characteristics. **B,** A minimal model with a single dimensionless variable $u$ for the signaling state of each cell (Eq. 1) captures the bistable behavior. The bifurcation diagram for cell state $u_1$ as a function of the signaling partner $u_2$ for symmetric coupling $c_{12} = c_{21} = 1$ shows the bistable regime, in which the two stable steady states (solid lines) are separated by an unstable state (dashed line). **C,** The signaling dipole of a pair of cells is given by Eq. 2 in the main text. For the arrangement shown here, the normalized axial component is given by $m = (u_2 - u_1)/2$ and is governed by the double-well potential Eq. 5 (Fig. 5A). In the absence of biasing factors, the positive and negative dipole configurations at $+1$ and $-1$ are degenerate. **D,** Mature hair-cell pairs almost always form positive dipoles. By imaging fluorescently labelled pairs of sibling hair cells over time, we found that 42 of 44 pairs were in a positive dipolar configuration during embryonic development, and 19 of 21 pairs had a positive dipole when regenerating after damage. The remaining pairs had undeterminable or uniform polarity; we never observed negative dipoles in mature cells. **E,** To describe how the dipole $\mathbf{m}$ changes orientation, we introduce the angle $\psi$ between $\mathbf{m}$ and the organ axis $x$.

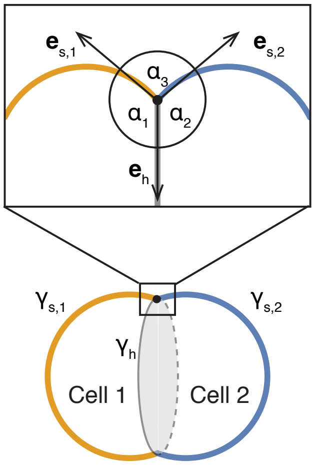


**Supplementary Figure 2: Geometric definitions at the contact point between the hair cells.** During the early stages of maturation, a pair of hair cells forms a doublet with a shared interface (denoted with subscript h) and interfaces between each cell and the surrounding supporting cells (subscripted $s, 1$ and $s, 2$). At the contact point between the cells, the angles $\alpha_1$, $\alpha_2$, and $\alpha_3$ between the three interfacial tangent vectors $\mathbf{e}_{s,1}$, $\mathbf{e}_{s,2}$ and $\mathbf{e}_h$ depend on the surface tensions $\gamma_h$, $\gamma_{s,1}$, and $\gamma_{s,2}$.

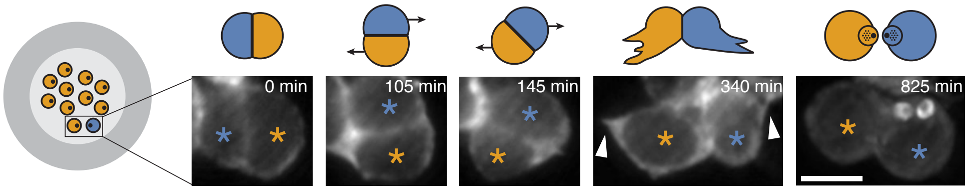


**Supplementary Figure 3: An oppositely polarized hair cell in a Notch mutant rearranges and extends protrusions.** Oppositely polarized hair cell pairs occasionally occurred in the neuromasts of Notch mutants. Fluorescent images from one such cell pair expressing $\beta$-actin-GFP show that the cells exchange positions along the PCP axis and subsequently form protrusions (arrowheads) similar to the oppositely polarized pairs of wild-type larvae. Scale bar: 5 µm.

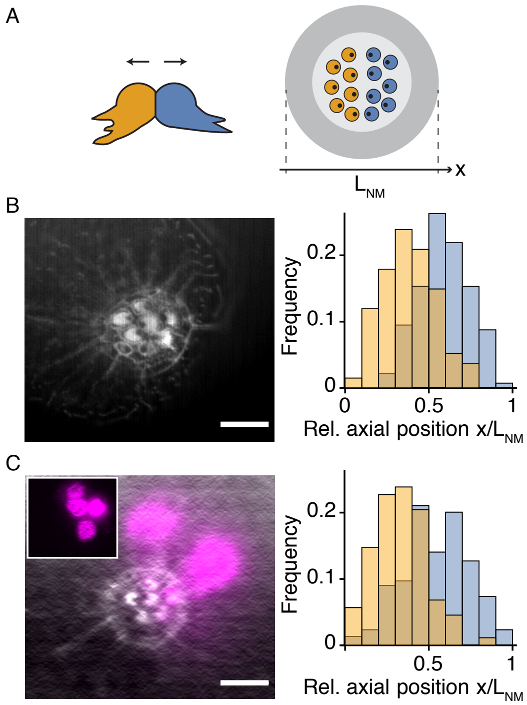


**Supplementary Figure 4: Polarity-directed positioning of hair cells.** **A,** In a schematic illustration of polarity-directed hair-cell positioning, oriented movements (left) sort the cells into the two halves of the organ, such that a mirror-symmetric polarity pattern appears (right). **B,** In wild-type neuromasts, hair cells of the two polarities (yellow and blue) differ significantly in their final axial position ($p < 0.0001$ for Mann Whitney test, 271 cells from 45 neuromasts). **C,** In larvae expressing constitutive NICD (magenta) under a stochastic hair cell-specific promotor, hair cells of the two polarities remain significantly different in their axial positions ($p < 0.0001$ for Mann Whitney test, 387 cells from 84 neuromasts). The datasets were obtained from images that were acquired as part of another study [1]. Scale bars: 5 µm.

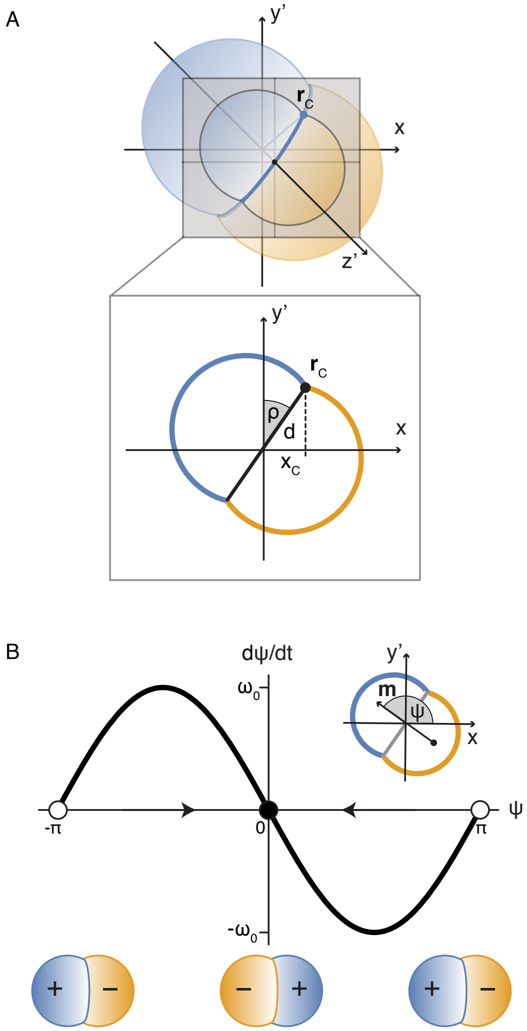


**Supplementary Figure 5: Mechanics of hair-cell rearrangements.** **A,** Active movements of the two hair cells along the external organ axis $x$ can lead to a rotation of the contact surface. A point on the contact line $\mathbf{r}_\mathrm{C} = (x_\mathrm{C}, y'_\mathrm{C}, z'_\mathrm{C})$ can be parameterized in the $x, y'$-plane by the distance $d$ to the axis of rotation $z'$ and the rotated angle $\rho$, such that $x_\mathrm{C} = d \sin \rho$. **B,** The velocity of the dipole angle $\psi$ is given by $-\omega_0 \sin \psi$. Any deviation from the unstable steady states (white circles), for which the cell pair forms a negative dipole relative to the organ axis, lead to angular movements that restore a positive axial dipolar state at the stable steady state (black circle).

### Supplementary Note

#### List of symbols

| | |
|---|---|
| $i, j...$ | Cell index |
| $u$ | Signaling state |
| $\tau$ | Characteristic time |
| $\sigma$ | Sigmoidal function |
| $s$ | Signal |
| $c$ | Coupling coefficient |
| $\mathbf{m}$ | Dipole |
| $\mathbf{r}$ | Position vector |
| $L$ | Cell-cell distance |
| $E$ | Potential |
| $\psi$ | Dipole angle |
| $(\mathrm{h}), (\mathrm{s}, 1), (\mathrm{s}, 2)$ | Surface subscripts |
| $\mathcal{A}$ | Cellular surface area |
| $\gamma$ | Surface tension |
| $\alpha$ | Contact angle |
| $f$ | Force per length |
| $\mathbf{e}$ | Unit tangent vector |
| $\delta$ | Perturbation |
| $\lambda$ | Length-scale |
| $\xi$ | Friction coefficient |
| $\mathbf{v}$ | Velocity |
| $d$ | Radial distance |
| $\rho$ | Rotated angle |
| $M$ | Torque |
| $\omega$ | Angular velocity |

### 1 Notch signaling dynamics of hair cell pairs

#### 1.1 Mutual inhibition in Notch signaling

The Notch signaling pathway regulates many processes in morphogenesis. Although biochemical signaling often relies on long-range transport of soluble ligand molecules by diffusion or active convection, both the Notch receptor and its ligands are bound to membranes on the cellular surfaces, such that cells engaged in Notch signaling must share a direct contact. Binding between receptors and ligands at intercellular contacts trigger intracellular processes that alter gene expression.

In simplified terms, signaling is initiated when a Notch-expressing cell binds an appropriate ligand on an adjacent cell (Supplementary Fig. 1A). In most contexts, the activation of the pathway downregulates the expression of ligand in the activated cell. Because the ability of a cell to *send* a signal is accordingly suppressed upon *receiving* a signal, Notch signaling gives rise to bistable cellular dynamics. Small initial differences in the expression levels of ligands and receptors between interacting cells are amplified in a winner-takes-all scenario, and cells with different signaling states subsequently differentiate into distinct cell types.

#### 1.2 A minimal model of Notch signaling

Here we introduce a minimal model of Notch signaling, modified from [2]. The signaling state of a cell is described by a single, unitless state variable $u$, whose dynamics depends on the signal $s$ received by that cell. For $N$ cells indexed $i \in 1, 2, ..., N$, the dynamical equation for a cell $i$ is given by

$$\tau \frac{du_i}{dt} = \sigma(u_i - s_i) - u_i, \tag{1}$$

with a timescale $\tau$ and a generic sigmoidal function $\sigma(u)$, here chosen as $\sigma(u) = \tanh(2u)$. The received signal $s_i$ is produced by surrounding cells, and in general depends on their respective signaling states. For our purposes, it suffices to consider $s$ to linear order in $u$, such that we may write

$$s_i = \sum_{j=1}^{N} c_{ij} u_j \tag{2}$$

with $c_{ij}$ the coefficient for the coupling between cells $i$ and $j$, in which we generally assume $c_{ii} = 0$. The system defined by Eq. 1 has two monostable regimes, separated by a bistable regime, with the two stable steady states corresponding to the ligand-expressing and receptor-expressing states respectively (Supplementary Fig. 1B).

#### 1.3 The signaling dipole

To describe the spatial arrangement of oppositely polarized cells within the neuromast, it is useful to introduce the signaling dipole moment $\mathbf{m}$ for a group of cells as

$$\mathbf{m} = \sum_{i=1}^{N} u_i \left( \mathbf{r}_i - \mathbf{r}_0 \right), \tag{3}$$

with $\mathbf{r}_i = (x_i, y_i, z_i)$ the position vector of cell $i$, and $\mathbf{r}_0$ the centroid of the collective configuration.

Only two cells are involved in the signaling interaction that establishes hair-cell polarity. For two cells of the same size, the pair dipole is given by Eq. 2 in the main text. The two cells arise from the division of a progenitor cell. The division axis is typically aligned with the external organ axis as a consequence of planar cell polarity signaling. Without loss of generality, we consider here the configuration shown in Supplementary Fig. 1C, in which the external polarity axis of the organ corresponds to the Cartesian x-axis. The axial component normalized by the cell-cell distance $L$ is then given by $m = (u_2 - u_1)/2$, for which we obtain the following dynamical equation using

Eqs. 1–2:

$$\tau \frac{dm}{dt} = \sigma(2m) - m, \tag{4}$$

in which we have set the couplings to $c_{12} = c_{21} = 1$. Equation 4 represents a bistable system with double-well potential (Fig. 5A):

$$E(m) = E_0 - \int (\sigma(2m) - m)\, dm = E_0 + \frac{m^2}{2} - \frac{\log(\cosh(4m))}{4}. \tag{5}$$

In the following sections, we consider rotations of the dipole $\mathbf{m}$, that result from cellular movements along the organ axis $x$. Without any additional broken symmetries, the axis of rotation is arbitrary, and we can parameterize the plane of rotation by $x$ and a perpendicular coordinate $y'$, such that the dipole expressed in the basis $\mathbf{e}_x, \mathbf{e}_{y'}$ reads

$$\mathbf{m} = uL(\cos\psi \mathbf{e}_x + \sin\psi \mathbf{e}_{y'}), \tag{6}$$

in which $\psi$ denotes the dipole angle between $\mathbf{m}$ and $x$, and $u = |u_i|$ (Supplementary Fig. 1E). Fig. 5A in the main text shows the three-dimensional potential $E(\mathbf{m})$.

### 2 Surface mechanics of hair-cell maturation

Neuromast hair cells undergo distinct rearrangements of their cellular interfaces over the course of their maturation. Here we introduce the theoretical framework used to calculate the predictions for cellular shape changes and movements (Figs. 2 and Fig. 4).

First, we describe the dynamics of the contact angle between the two hair cells in terms of the underlying changes in their effective interfacial tensions. This process takes place over the entire duration of hair-cell maturation, and is likely driven by gradual changes in gene expression as the two cells differentiate into hair cells.

Next, we consider the mechanical consequences of the biochemical breaking of symmetry described in Supplementary Note 1. We explore the effects of oppositely oriented cell-intrinsic spatial gradients in surface tension in the two cells, in which the sign of the gradient relative to the organ axis is set by $u_i$ in each cell. As supported by our experimental results, we assume that the two cells are otherwise identical, and in particular that they have the same volume. See also [1] for experimental findings that argue against the possibility that the hair-cell progenitor undergoes an asymmetric cell division.

#### 2.1 Force balance at the contact line

After the division of a precursor cell, the two sibling hair cells adhere to one another and are otherwise surrounded by supporting cells. Each hair cell thus has two distinct types of contact surfaces, and the line of contact points between the hair cells and the supporting cells is formed by the boundaries of three interfaces. Let $\mathbf{r}_\mathrm{C}$ denote a point on the contact line relative to $\mathbf{r}_0$, and the subscript (h) label all variables and parameters associated with the interface between the two hair cells, and the subscripts $(\mathrm{s}, i)$ those associated with the interface of cell $i$ and the supporting cells. Furthermore, we denote the angles between the interfaces by $\alpha_1$, $\alpha_2$, and $\alpha_3$ respectively (Supplementary Fig. 2). The interfacial areas $\mathcal{A}_\mathrm{h}$, $\mathcal{A}_{\mathrm{s},1}$, and $\mathcal{A}_{\mathrm{s},2}$ are dual to the interfacial tensions $\gamma_\mathrm{h}$, $\gamma_{\mathrm{s},1}$, and $\gamma_{\mathrm{s},2}$, which give rise to so-called Young forces that act on the point $\mathbf{r}_\mathrm{C}$ ([3]). The total Young force per unit length can be written as $\mathbf{F}_\mathrm{Y} = f_\mathrm{h}\mathbf{e}_\mathrm{h} + f_{\mathrm{s},1}\mathbf{e}_{\mathrm{s},1} + f_{\mathrm{s},2}\mathbf{e}_{\mathrm{s},2}$, in which $\mathbf{e}_\mathrm{h}$, $\mathbf{e}_{\mathrm{s},1}$, and $\mathbf{e}_{\mathrm{s},2}$ denote the unit tangent vectors of the respective interfaces defined normal to the contact line (Supplementary Fig. 2). The components of $\mathbf{F}_\mathrm{Y}$ in the direction along each of the interfaces are given by

$$f_{\mathrm{h}} = \gamma_{\mathrm{h}} + \gamma_{\mathrm{s},1} \cos \alpha_1 + \gamma_{\mathrm{s},2} \cos \alpha_2, \tag{7}$$

$$f_{\mathrm{s},1} = \gamma_{\mathrm{s},1} + \gamma_{\mathrm{s},2} \cos \alpha_3 + \gamma_{\mathrm{h}} \cos \alpha_1, \tag{8}$$

$$f_{\mathrm{s},2} = \gamma_{\mathrm{s},2} + \gamma_{\mathrm{h}} \cos \alpha_2 + \gamma_{\mathrm{s},1} \cos \alpha_3. \tag{9}$$

When the system is in a static equilibrium, no net forces arising from the interfacial tensions act on the contact point, i.e. $\mathbf{F}_{\mathrm{Y}} = \mathbf{0}$. For $\gamma_{\mathrm{s},1} = \gamma_{\mathrm{s},2} = \gamma_{\mathrm{s}}$, and using the geometric constraint $\alpha_1 + \alpha_2 + \alpha_3 = 2\pi$, we then obtain the following relation for the equilibrium contact angle $\alpha_3^{\mathrm{E}}$

$$\cos\left(\frac{\alpha_3^{\mathrm{E}}}{2}\right) = \frac{\gamma_{\mathrm{h}}}{2\gamma_{\mathrm{s}}}, \tag{10}$$

which corresponds to Eq. 4 in the main text with the choice $\alpha = \alpha_3^{\mathrm{E}}/2$ [4].

#### 2.2 Contact-angle dynamics during hair-cell maturation

We observed that the hair-cell contact angle $\alpha$ decreases over the course of maturation in a highly stereotyped manner, concluding in the detachment of the two cells from each other. Based on evidence of differential adhesion between hair cells and supporting cells in the mammalian cochlea [5, 6], we hypothesize that the dynamics of the contact angle is determined primarily by an increase in the heterophilic adhesion at the interfaces between the hair and supporting cells. Assuming that hair-cell differentiation leads to a gradual change in $\gamma_{\mathrm{s}}$ over a time interval $t \in [0, T]$, with $T$ denoting the timepoint of cell-cell detachment, we consider the series expansion of $\gamma_{\mathrm{s}}$ to first order

$$\gamma_{\mathrm{s}}(t/T) = \gamma_0 + \gamma_1 t/T + \mathcal{O}((t/T)^2). \tag{11}$$

At the time of detachment, the contact angle is zero, and from Eq. 10 it therefore follows that $\gamma_\mathrm{h}(T) = 2\gamma_\mathrm{s}(T)$. Assuming that the surface interactions between the two hair cells remain unchanged so that $\gamma_\mathrm{h}$ remains constant, we obtain from Eq. 11 the following expression for the tension ratio over time

$$\cos\alpha = \frac{\gamma_\mathrm{h}}{2\gamma_\mathrm{s}} = \frac{1+\tilde{\gamma}}{1+\tilde{\gamma}t/T}, \tag{12}$$

with $\tilde{\gamma} = \gamma_1/\gamma_0$. Fitting $\tilde{\gamma}$ to the experimental measurements by nonlinear least-squares minimization yields the estimate $\tilde{\gamma} = -0.93 \pm 0.01$.

In using the equilibrium relation here, we assume that $\gamma_\mathrm{s}$ changes slowly compared to the relaxation of the contact angle. Indeed, the timescale of the tension ratio is on the order of hours (Fig. 2), whereas any slight shape fluctuations we observed occurred on a minute scale. A fraction of the hair-cell pairs also undergoes a dynamical rearrangement process on a timescale of tens of minutes that is neglected here (Fig. 4). We describe the details of this fast rearrangement process in Supplementary Note 2.4, in which we rely self-consistently on the converse timescale-separation argument and take $\gamma_\mathrm{s}$ to be quasi-static.

#### 2.3 Asymmetric supporting-cell tensions

In this section, we study the forces arising from an asymmetry between the two cells in the tensions $\gamma_{\mathrm{s},i}$. Focusing on a timescale that is small compared to the characteristic timescale of the contact-angle dynamics, we examine the initial period of hair-cell maturation, such that we can take the surface tensions to be stationary over the relevant time interval and we can consider small perturbations from a symmetric state of the form $\gamma_{\mathrm{s},i} = \gamma_\mathrm{s} + \delta\gamma_{\mathrm{s},i}$ (Fig. 4A). For parameter regimes in which the angles are differentiable functions of the tensions, we may write

$$\alpha_i = \pi - \cos^{-1}\left(\frac{\gamma_\mathrm{h}}{2\gamma_\mathrm{s}}\right) + \sum_{j=1}^{2} \left.\frac{\mathrm{d}\alpha_i}{\mathrm{d}\delta\gamma_{\mathrm{s},j}}\right|_{\delta\gamma_{\mathrm{s},j}=0} \delta\gamma_{\mathrm{s},j} + \mathcal{O}(\delta^2), \tag{13}$$

$$\alpha_3 = 2\cos^{-1}\left(\frac{\gamma_\mathrm{h}}{2\gamma_\mathrm{s}}\right) + \sum_{i=1}^{2} \left.\frac{\mathrm{d}\alpha_3}{\mathrm{d}\delta\gamma_{\mathrm{s},i}}\right|_{\delta\gamma_{\mathrm{s},i}=0} \delta\gamma_{\mathrm{s},i} + \mathcal{O}(\delta^2), \tag{14}$$

and expand the forces to linear order in the perturbations $\delta\gamma_{\mathrm{s},i}$, obtaining

$$f_\mathrm{h} = -\frac{\gamma_\mathrm{s}}{2}\sqrt{4 - \frac{\gamma_\mathrm{h}^2}{\gamma_\mathrm{s}^2}}(\delta\alpha_1 + \delta\alpha_2) - \frac{\gamma_\mathrm{h}(\delta\gamma_{\mathrm{s},1} + \delta\gamma_{\mathrm{s},2})}{2\gamma_\mathrm{s}} + \mathcal{O}(\delta^2), \tag{15}$$

$$f_{\mathrm{s},1} = -\frac{\gamma_\mathrm{h}}{2}\sqrt{4 - \frac{\gamma_\mathrm{h}^2}{\gamma_\mathrm{s}^2}}\delta\alpha_1 - \gamma_\mathrm{s}\sin\left(2\sec^{-1}\left(\frac{2\gamma_\mathrm{s}}{\gamma_\mathrm{h}}\right)\right)\delta\alpha_3 + \cos\left(2\sec^{-1}\left(\frac{2\gamma_\mathrm{s}}{\gamma_\mathrm{h}}\right)\right)\delta\gamma_{\mathrm{s},2} + \delta\gamma_{\mathrm{s},1} + \mathcal{O}(\delta^2), \tag{16}$$

$$f_{\mathrm{s},2} = -\frac{\gamma_\mathrm{h}}{2}\sqrt{4 - \frac{\gamma_\mathrm{h}^2}{\gamma_\mathrm{s}^2}}\delta\alpha_2 - \gamma_\mathrm{s}\sin\left(2\sec^{-1}\left(\frac{2\gamma_\mathrm{s}}{\gamma_\mathrm{h}}\right)\right)\delta\alpha_3 + \cos\left(2\sec^{-1}\left(\frac{2\gamma_\mathrm{s}}{\gamma_\mathrm{h}}\right)\right)\delta\gamma_{\mathrm{s},1} + \delta\gamma_{\mathrm{s},2} + \mathcal{O}(\delta^2), \tag{17}$$

in which the angle perturbations are abbreviated as $\delta\alpha$. At mechanical equilibrium, Eqs. 15–17 are solved by

$$\delta\alpha_1^\mathrm{E} = \frac{2\gamma_\mathrm{s}^2(\delta\gamma_{\mathrm{s},1} - \delta\gamma_{\mathrm{s},2}) - \gamma_\mathrm{h}^2\delta\gamma_{\mathrm{s},1}}{\gamma_\mathrm{h}\gamma_\mathrm{s}^2\sqrt{4 - \frac{\gamma_\mathrm{h}^2}{\gamma_\mathrm{s}^2}}}, \tag{18}$$

$$\delta\alpha_2^\mathrm{E} = \frac{2\gamma_\mathrm{s}^2(\delta\gamma_{\mathrm{s},2} - \delta\gamma_{\mathrm{s},1}) - \gamma_\mathrm{h}^2\delta\gamma_{\mathrm{s},2}}{\gamma_\mathrm{h}\gamma_\mathrm{s}^2\sqrt{4 - \frac{\gamma_\mathrm{h}^2}{\gamma_\mathrm{s}^2}}}, \tag{19}$$

$$\delta\alpha_3^\mathrm{E} = \frac{\gamma_\mathrm{h}(\delta\gamma_{\mathrm{s},1} + \delta\gamma_{\mathrm{s},2})}{\gamma_\mathrm{s}^2\sqrt{4 - \frac{\gamma_\mathrm{h}^2}{\gamma_\mathrm{s}^2}}}. \tag{20}$$

Away from the static equilibrium configuration, however, unbalanced Young forces act on the contact line and give rise to dissipative restoring forces [3, 7]. It is illustrative to consider the limit $\gamma_{\mathrm{h}} \to 0$, for which the equilibrium solution ceases to exist altogether. In this limit, the angle $\alpha_3$ approaches $\pi$ whereas Eqs. 18–19 diverge to infinity. Asymmetric and uniform supporting cell interfacial tensions $\gamma_{\mathrm{s},i}$ lead to a final configuration in which the contact line described by Eqs. 8–7 is lost. For example, if $\gamma_{\mathrm{s},1} > \gamma_{\mathrm{s},2}$, cell 2 eventually internalizes cell 1 [8], because the difference in surface tension gives rise to a displacement of the contact line, which increases the size of the interface with the lower surface tension at the expense of the interface with the higher surface tension. As long as the system is out of static equilibrium, uncompensated Young forces are balanced by frictional forces that depend on the velocity of the contact line. Each point along the contact line moves at a uniform velocity that depends on the tension difference $\gamma_{\mathrm{s},1} - \gamma_{\mathrm{s},2}$ acting on the line, such that the contact line preserves its circular shape and shrinks in size until it becomes a point at the pole and finally disappears.

Non-uniform surface tensions however can give rise to stable Galilean-invariant moving solutions, in which the angles Eqs. 13–14 take on dynamic steady-state values (sometimes called "apparent" contact angles) instead of approaching a singularity. Such problems have been studied in various contexts, including for example droplets moving on flat substrates with chemical adhesion gradients [7] or cells with active surface tension gradients moving in cylindrical pipes [9]. In the next section, we explore the effect of non-uniform surface tensions in the hair-cell doublet.

#### 2.4 Gradient in surface tension

We next consider the case in which the surface tensions are modulated by a cell-intrinsic polarity field. The signaling state $u_i$ of cell $i$ is determined by biochemical interactions (see Supplementary Note 1), and as before, the Cartesian x-axis is defined as the external axis of the organ. The two hair cells are assumed to be identical at the outset, but as they undergo Notch signaling across their contact surface, they establish their respective polarity identities, and the perturbations $\delta\gamma_{s,1}$ and $\delta\gamma_{s,2}$ become non-zero as well as spatially non-uniform. The cell-intrinsic polarities are then oriented in opposite directions along the polarity axis $x$, with their sign determined by the steady-state values of the respective signaling states. We consider a small, linear gradient, and define the perturbations as

$$\delta\gamma_{s,i} = -\frac{u_i \Delta\gamma_s}{\lambda}(x - x_i), \tag{21}$$

in which $\Delta\gamma_s$ is the magnitude of the tension drop along the polarity axis, $x_i$ denotes a reference point within the cell, and $\lambda$ is a gradient length scale on the order of the cell size. Note that for the following calculations, the cell-intrinsic reference point of the tension kernel $x_i$ can be chosen arbitrarily, but with the choice $x_i = x_0$, in which $x_0$ is the centroid of the cell pair, our measurement of the contact angle over the maturation time can be used self-consistently to estimate the slow dynamics of the baseline tension $\gamma_s$ (Fig. 2; Supplementary Note 2.2), which we used in constructing the state diagram Fig. 4A.

We furthermore assume that $\gamma_h$ is finite but small (i.e. of order $\delta\gamma_{s,i}$), which substantially simplifies the problem. In the symmetric reference state, the angles $\alpha_3$ and $\alpha_{1,2}$ then have equilibrium solutions close to $\pi$ and $\pi/2$ respectively, and the dynamic angle perturbations can be assumed to remain small and uniform as long as intracellular pressure differences equilibrate on timescales that are fast compared to any movement of the contact line (see also [7]). By symmetry, it then follows that the contact line is circular and the contact surface between the hair cells remains without curvature. Indeed, we observe that the shape of the cell doublet is nearly spherical over the relevant time period with angles close to $\alpha_3 = \pi$ and $\alpha_{1,2} = \pi/2$ (Figs. 2 and 4; Supplementary Video 2). With these assumptions, the linearized

Young forces simplify further to

$$f_{\mathrm{h}} = -\gamma_{\mathrm{s}}(\delta\alpha_1 + \delta\alpha_2) + \mathcal{O}(\delta^2), \tag{22}$$

$$f_{\mathrm{s},1} = \delta\gamma_{\mathrm{s},1} - \delta\gamma_{\mathrm{s},2} + \mathcal{O}(\delta^2), \tag{23}$$

$$f_{\mathrm{s},2} = -(\delta\gamma_{\mathrm{s},1} - \delta\gamma_{\mathrm{s},2}) + \mathcal{O}(\delta^2). \tag{24}$$

From Eqs. 23–24 it follows that there is no static solution for an equilibrium angle, and the uncompensated Young force must be balanced by a drag force $\mathbf{F}_{\mathrm{D}}$ at each point along the contact line. In general, the dissipative force depends on the velocity of the contact point, and for small $\mathbf{v}$ we may use here a linear relationship $\mathbf{F}_{\mathrm{D}} = \xi\mathbf{v}$, with $\xi$ a generic dissipative coefficient. Conservation of momentum requires $\mathbf{F}_{\mathrm{Y}} = \mathbf{F}_{\mathrm{D}}$, from which we obtain $\mathbf{v} = \mathbf{F}_{\mathrm{Y}}/\xi$ for the velocity at each point of the contact line.

It is helpful to decompose the velocity into a pair of components, one parallel and one perpendicular to $\mathbf{r}_{\mathrm{C}}$, such that $\xi\mathbf{v}_\perp = \sum_i \sin\alpha_i f_{\mathrm{s},i}\mathbf{e}_{\mathrm{s},i}$ and $\xi\mathbf{v}_\parallel = f_{\mathrm{h}}\mathbf{e}_{\mathrm{h}} + \sum_i \cos\alpha_i f_{\mathrm{s},i}\mathbf{e}_{\mathrm{s},i}$. These equations describe the radial and angular motion of the point $\mathbf{r}_{\mathrm{C}}$. To linear order, we find that the parallel velocity depends only on the angles, and the perpendicular velocity only on the tension perturbations,

$$v_\parallel = |\mathbf{v}_\parallel| = \frac{\gamma_{\mathrm{s}}(\delta\alpha_1 + \delta\alpha_2)}{\xi} + \mathcal{O}(\delta^2), \tag{25}$$

$$v_\perp = |\mathbf{v}_\perp| = \frac{4u\Delta\gamma_{\mathrm{s}}}{\lambda\xi} x_{\mathrm{C}} + \mathcal{O}(\delta^2), \tag{26}$$

in which we have substituted the tension perturbations (Eq. 21), and used $u = |u_1| = |u_2|$ as in Eq. 6. Together with the constraint that the two cell volumes are conserved, the radial equation Eq. 25 can be used to compute the dynamic angles, and Eq. 26 yields the angular velocity $\omega = |\mathbf{v}_\perp|/d$, in which $d$ is the distance of $\mathbf{r}_{\mathrm{C}}$ from the axis of rotation. For the negative dipolar configuration shown in Supplementary Fig. 5A, we can substitute

$x_C = d \sin \rho$, in which $\rho$ is the rotated angle of $\mathbf{r}_C$, and find

$$\omega = \omega_0 \sin \rho + \mathcal{O}(\delta^2), \tag{27}$$

with $\omega_0 = 4u\Delta\gamma_s/(\lambda\xi)$. Because $\omega_0$ does not depend on the position along the contact line and is constant in $\rho$, it follows that the movement of the contact line can be described for small gradients as a rigid-body rotation.

The angular velocity of the dipole $\mathbf{m}$ can be obtained from Eq. 27 through the relation between the rotated angle $\rho$ and the dipole angle $\psi$ as defined in Eq. 6, and is given by $\mathrm{d}\psi/\mathrm{d}t = -\omega_0 \sin \psi$. Thus, for the initial states in which $\psi = \pm\pi$, any small fluctuation in $\psi$ triggers a rotation of the interface by $\pi$ until the stable configuration at $\psi = 0$ is achieved. Conversely, if the pair is initially in a positive dipolar configuration, fluctuation-induced angular movements act to restore the dipole angle to $\psi = 0$ (Supplementary Fig. 5B).

The magnitude of the angular velocity depends on the magnitude of the surface tension drop $\Delta\gamma_s$.

##### 2.4.1 Surface energy of the cell doublet

Finally, we present another derivation of the rotational velocity of the interface between the sibling hair cells that is based on energetic considerations. The total effective surface energy of the cell doublet is given by Eq. 3 in the main text. For simplicity, the shape of the cell doublet is taken as spherical with radius $R$ in the following. Note that our experimental observations of rotating cell doublets confirm that deviations from the spherical shape are small (Fig. 4B), in agreement with our assumption that the tension gradients are small during the relevant time interval.

For the configuration in Supplementary Fig. 5A, we consider a reference frame rotated by $\rho$, parameterize the co-rotated cell-doublet surface with spherical coordinates $\theta$ and $\phi$ and calculate the transformed surface tensions

Eq. 21, obtaining

$$\gamma_{\mathrm{s},i} = \gamma_{\mathrm{s}} - u_i \frac{R\Delta\gamma_{\mathrm{s}}}{\lambda}(\cos\phi\sin\theta\sin\rho - \cos\theta\cos\rho). \tag{28}$$

Substituting Eq. 28 into the expression for the surface potential (Eq. 3), and evaluating the integrals over the hemispherical surfaces, we obtain

$$\frac{E - \mathcal{A}_{\mathrm{h}}\gamma_{\mathrm{h}}}{(\mathcal{A}_{\mathrm{s},1} + \mathcal{A}_{\mathrm{s},2})\gamma_s} = 1 + u\frac{R}{\lambda}\frac{\Delta\gamma_s}{2\gamma_s}\cos\rho. \tag{29}$$

Equation 7 in the main text corresponds to Eq. 29 expressed as a function of the dipole angle $\psi$ and with the simplification $R/\lambda \approx 1$. Correspondingly, the unstable equilibrium position of Eq. 29 is at $\rho = 0$ and the stable position at $\rho = \pi$. The derivative of the surface potential with respect to $\rho$ defines a torque that is related to the angular velocity through an angular friction coefficient $\xi_\omega$ as

$$\omega = -\frac{1}{\xi_\omega}\frac{\partial E}{\partial \rho} = \omega_0 \sin\rho, \tag{30}$$

in agreement with our previous result (Eq. 27). Comparison of the theoretically calculated angular velocity and the measurements of cell doublets lead to a fit parameter $\omega_0 = (0.25 \pm 0.01)10^{-3}{\cdot}\pi/s$ (Fig. 4E). Although we calculate $\omega$ under the assumption that the surface tensions are constant over the relevant time interval, our results indicate changes in both $\Delta\gamma_{\mathrm{s}}$ and $\gamma_{\mathrm{s}}$ over long timescales (Figs. 2 and 4A), which could be responsible for the slight underestimation of the angular velocity in the right branch of the profile.

Note that our observations did not suggest any additional broken symmetries, such as a preferred axis of rotation. Accordingly, in our description, the breaking of symmetry occurs with respect to the external organ axis $x$ only, whereas the other two axes $z'$ and $y'$, which form the Cartesian reference frame of the pair, are chosen by the initial fluctuation of the interface and are therefore allowed to vary from pair to pair across a neuromast.
